# Role of the *EGR2* promoter antisense RNA in modulating chromatin accessibility and spatial genome organization

**DOI:** 10.1101/2022.05.02.490330

**Authors:** Margot Martinez-Moreno, David Karambizi, Hyeyeon Hwang, Kristen Fregoso, Jia-Shu Chen, Eduardo Fajardo, Andras Fiser, Nikos Tapinos

## Abstract

The *EGR2* promoter antisense RNA (AS-RNA) recruits chromatin remodeling complexes to inhibit *EGR2* transcription following peripheral nerve injury. Here we show that the *EGR2*-AS-RNA modulates chromatin accessibility and interacts with two distinct histone modification complexes. It binds to EZH2 and WDR5 and enables targeting of H3K27me3 and H3K4me3 to promoters of *EGR2*and *C-JUN* respectively. Expression of the AS-RNA results in reorganization of the global chromatin landscape and quantitative changes in loop formation and in contact frequency at domain boundaries exhibiting enrichment for AP-1 genes. In addition, the *EGR2*-AS-RNA induces changes in hierarchical TADs and increases transcription factor occupancy on an inter-TAD loop between a super-enhancer regulatory hub and the promoter of *mTOR*. Our results show that the *EGR2*-AS-RNA may serve as regulator of chromatin remodeling and spatial genome organization in Schwann cells.

## Introduction

Spatial genome organization involves the three-dimensional (3D) structure, positioning, and interactions of chromatin within the nucleus. This is a non-random process and is prone to regulation within various nuclear domains, topological associations and through chromatin modifying epigenetic mechanisms. In recent years, non-coding RNAs (ncRNAs) have emerged as major regulators of spatial genome organization. ncRNAs are RNA molecules that are not translated into proteins and are implicated in numerous cellular processes including transcription, mRNA splicing, and protein translation ^1–3^. ncRNAs impact spatial genome organization by modulating perinuclear chromosome tethering, the formation of major nuclear compartments, chromatin looping and various chromosomal structures ^4^. These roles of ncRNAs often intersect with various other protein or nucleic acid components of genome structure and function. Although the role of ncRNAs as modulators of the 3D genome organization of the X Chromosome, various developmental genes and repetitive DNA loci ^4^ has been described, the exact mechanistic role of the low-copy promoter associated antisense RNAs (AS-RNAs) in gene regulation, chromatin remodeling and spatial genome organization is not known.

We have previously mapped an antisense RNA complementary to the *EGR2* promoter in Schwann cells (SCs), showed that its expression is increased following peripheral nerve injury and that it mediates the recruitment of EZH2 and H3K27me3 on *EGR2* promoter to induce transcriptional silencing of *EGR2* ^5^. Here we reveal the epigenetic function of this promoter AS-RNA (*EGR2*-AS-RNA) and show that it works as a molecular scaffold to bring WDR5, EZH2 together with activating (H3K4me3) and repressing (H3K27me3) histone marks on the promoters of *C-JUN* and *EGR2* respectively. Expression of the *EGR2*-AS-RNA increases chromatin accessibility in promoters with footprints of the AP1 transcription factor family. Finally, interrogation of 3D genome architecture using Hi-C reveals that expression of the *EGR2*-AS-RNA in SCs induces re-organization of the 3D architecture, quantitative changes in hierarchical topologically associating domains (TADs) and increased transcriptional regulation on an inter-TAD interaction between a super-enhancer regulatory hub and the promoter of *mTOR*. We propose a new model for promoter AS-RNA-mediated transcriptional regulation through recruitment of chromatin modulators to reorganize chromatin and 3D genome architecture in hubs that define cellular plasticity.

## Results

### The *EGR2*-AS-RNA interacts with EZH2 and WDR5 to enable targeting of H3K27me3 and H3K4me3 at *EGR2* and *C-JUN* promoters

We have shown that the *EGR2*-AS-RNA has a direct repressive effect on the *EGR2* promoter after sciatic nerve injury by recruiting components of the Polycomb Repressive Complex (PRC) ^5^. The repair SC phenotype is characterized by the activation of certain factors, such as C-JUN, and histone modifications are key to the regulation of these genes after injury ^6^. To determine whether the regulation of *EGR2* and *C-JUN* is mediated by the *EGR2*-AS-RNA in conjunction with chromatin modifying enzymes (CMEs) and histone marks, we performed RNA immunoprecipitation (RIP) and chromatin immunoprecipitation (ChIP) in SCs after overexpression of the *EGR2*-AS-RNA. We performed RIPs with antibodies against EZH2 and WDR5, and ChIPs with antibodies against the tri-methylated histone 27 (H3K27me3, repressive) and the tri-methylated histone 4 (H3K4me3, activating) marks followed by qPCR to detect either the presence of the AS-RNA following RIP, or the presence of *EGR2* and *C-JUN* promoters binding to the histone marks (ChIP). The RIP experiment shows that both EZH2 and WDR5 pull down the *EGR2*-AS-RNA compared to the control samples (Figure 1A). In addition, overexpression of the *EGR2*-AS-RNA in SCs results in significant increase of H3K27me3 binding to the *EGR2* promoter, and significant increase in association of H3K4me3 to the *C-JUN* promoter (Figure 1B). This interaction was reverted with the addition of specific inhibitory oligonucleotide GapMers that target the AS-RNA, which suggests that the interaction of H3K27me3 and H3K4me3 with *EGR2* and *C-JUN* respectively depends on the instructive role of the AS-RNA.

**Figure 1:**
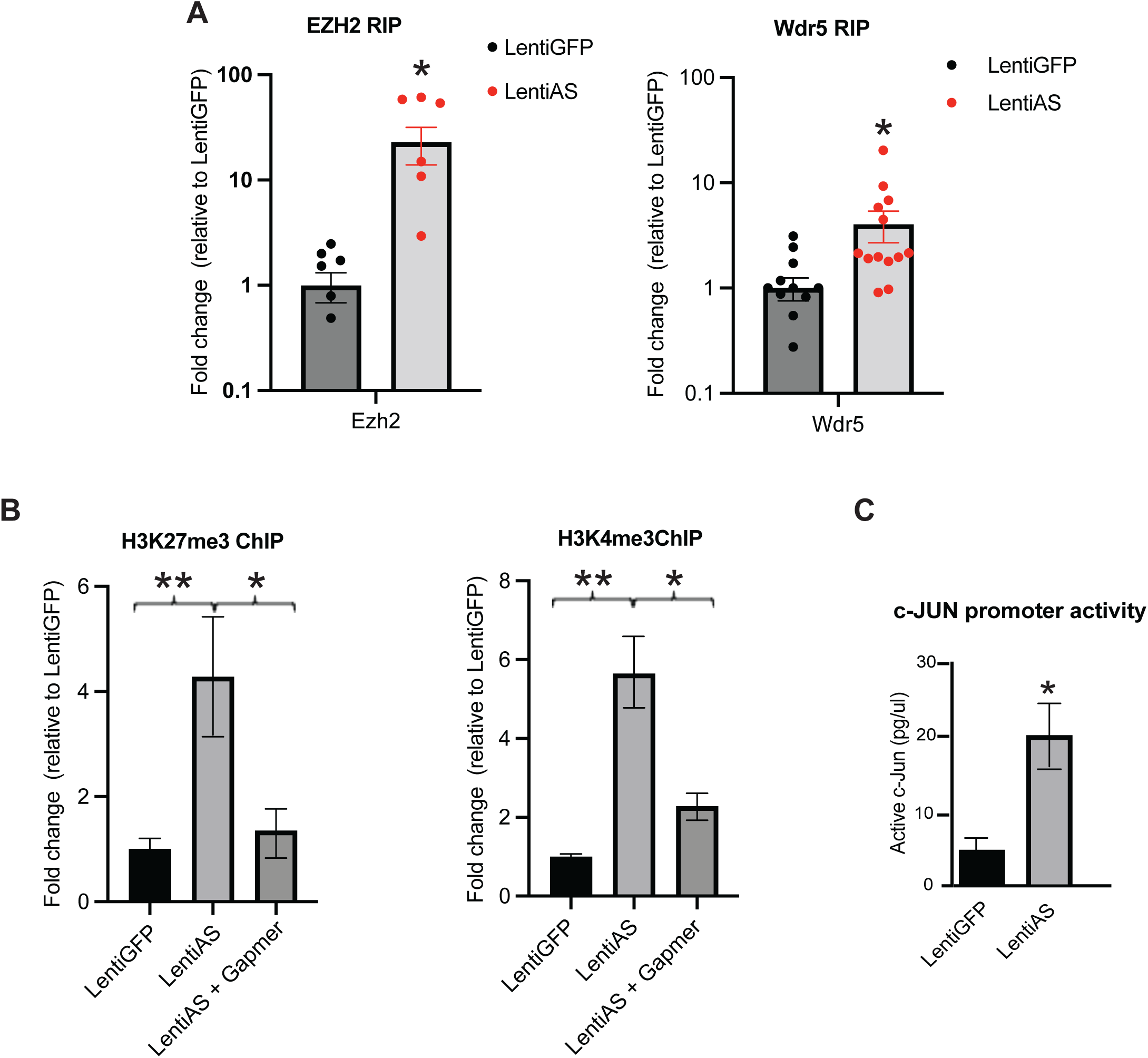
The *EGR2*-AS-RNA recruits EZH2 and WDR5 and enables targeting of H3K27me3 and H3K4me3 on *EGR2* and *C-JUN* promoters. A) RIP experiments after the *EGR2*-AS-RNA expression in SCs, with antibodies against EZH2 and WDR5. Significance was calculated with a Student’s t-test (For the WDR5 RIP, N = 13, 5 biological replicates, *p = 0.040, dF = 27; For the EZH2 RIP, N = 9, 3 biological replicates, *p = 0.026, dF = 16). B) ChIP experiments following expression of the EGR2 promoter AS-RNA in SCs and its effect to H3K27me3 binding on *EGR2* promoter and H3K4me3 binding on C-JUN promoter, respectively. Incubation of cells with oligonucleotide GapmeRs against the EGR2-AS-RNA, inhibits the AS-RNA induced binding of H3K27me3 and H3K4me3 on the EGR2 and C-JUN promoters. For the H3K27me3 ChIP, N = 13, 5 biological replicates and 1 technical replicate, *p = 0.020, dF = 23, **p = 0.0094, dF = 22. For the H3K4me3 ChIP, N = 15, 5 biological replicates and 1 technical replicate, *p = 0.049, dF = 18, **p = 0.0012, dF = 23. C) Overexpression of the *EGR2*-AS-RNA increases the C-JUN DNA binding activity compared to the control (N = 5/group, 3 independent experiments, *p = 0.0176, dF = 10).

To verify that expression of the AS-RNA has functional effects on promoter activity, we performed transcription factor (TF) activity assays following overexpression of the *EGR2*-AS-RNA to determine the activation of the transcription factor activity of C-JUN on the AP-1 specific binding motif. We show that the AS-RNA significantly increases TF activity of C-JUN (Figure 1C).

### The *EGR2*-AS-RNA induces chromatin remodeling and increased binding of the AP-1/JUN family of TFs

We performed ATAC-seq to capture genome wide epigenetic profile changes induced by the EGR2 promoter AS-RNA. First, we performed a PCA analysis, which showed that control and treated samples closely cluster together (Figure 2A, Figure S1A) suggesting that the epigenetic changes induced by the AS-RNA are reproducible. We then performed a differential accessibility analysis and observed that the AS-RNA increased accessibility at 109 and decreased it at 467 sites (Figure 2B). Genes with increased accessibility showed significant enrichment for the AP-1 transcription factor network (Figure 2C). Promoter of genes such as *JUN*, *JUNB*, *JUND*, *FOSB*, *FOS* saw a significant increase in accessibility following AS-RNA overexpression (Figure S1B).

**Figure 2:**
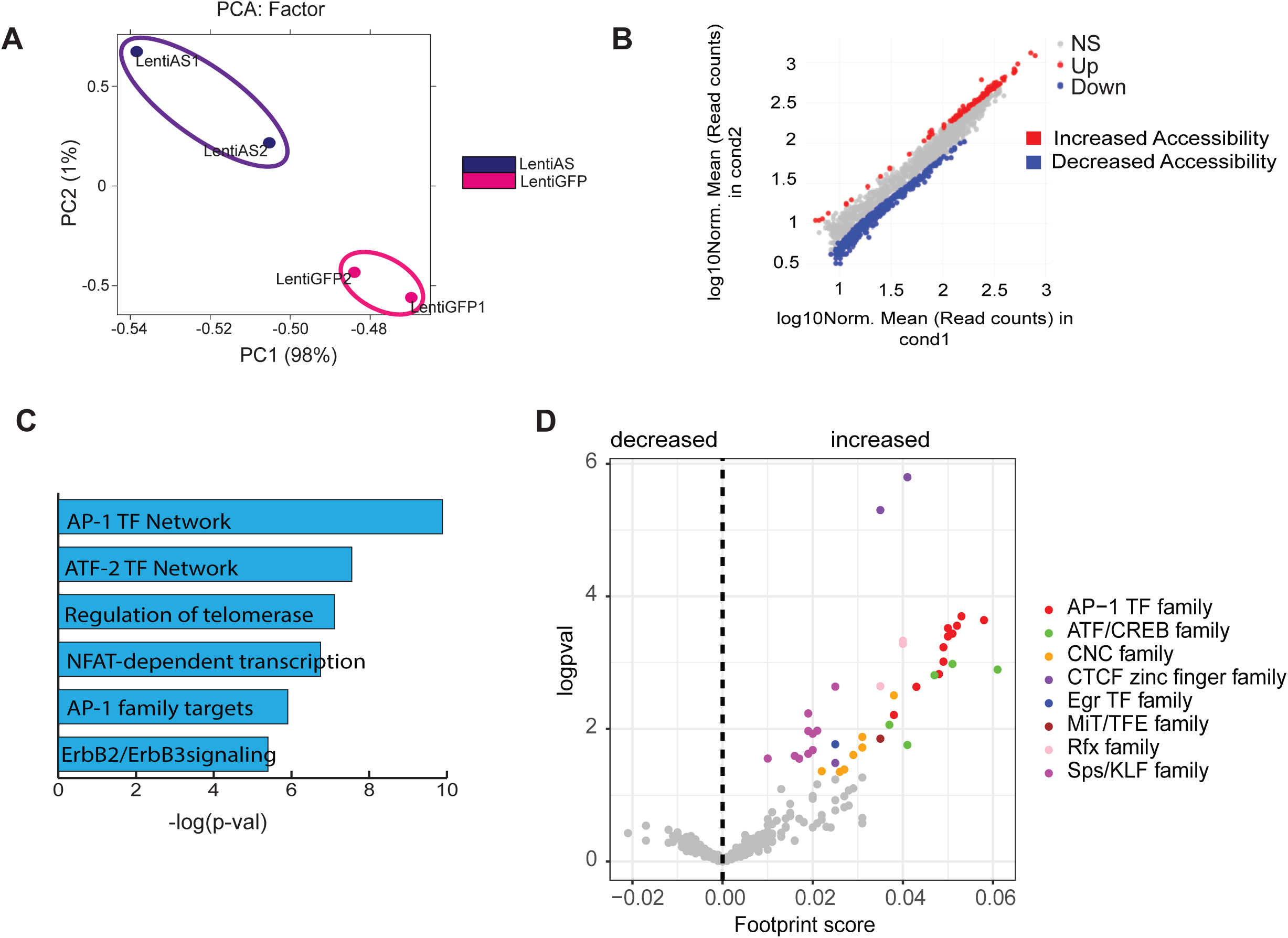
Expression of the *EGR2*-AS-RNA induces chromatin remodeling and increased binding of the AP-1/JUN TF family. A) Sample correlation analysis based on ATAC-seq peak location and intensity. PCA result of all samples is plotted as a 2D graph with PC1 as X-axis and PC 2 as Y-axis. Note that variable 1 is strong (98%) to divide into the 2 hierarchical groups. B) Scatterplot comparing ATAC-seq signal intensities across all open chromatin sites between the LentiAS group compared to the LentiGFP group. Significant changes correspond to an FDR-adjusted P value below 0.05 and an absolute log2 fold change above 1.5. The diagonal is shown as a gray area, as a reference indicating regions with no change in chromatin accessibility. Colored dots indicated differences in accessibility. C) GO analysis among genes located in the vicinity of regions with increased chromatin accessibility after the *EGR2*-AS-RNA overexpression. D) Volcano plot of differential TF binding shown as footprint score versus the significance (logpval) reveals that expression of the *EGR2*-AS-RNA induces significant increase in DNA binding of AP-1, CTCF, ATF/CREB, CNC, EGR, MiT/FTF, Rfx and KLF TF families.

Next, using accessibility data, we performed transcription factor (TF) footprinting analysis, which allows the prediction of the precise binding location of a TF at a particular locus. This is because the DNA bases that are directly bound by the TF are protected from transposition while the DNA bases immediately adjacent to TF binding are accessible. We found that the AP-1 TF family and the architectural protein CTCF (CTCF zinc finger family) experience the most significant increase in binding following AS-RNA expression (Figure 2D).

### Expression of the *EGR2*-AS-RNA results in genome reorganization and a gain in stable loops associated with genes enriched for AP-1 TF network and NOTCH 1 signaling

We hypothesized that expression of the Egr2-AS-RNA may induce spatial genome reorganization in SCs to epigenetically affect global transcriptional programs. We performed Hi-C following expression of the AS-RNA using Lenti-AS and compared the genome architecture with SCs expressing Lenti-GFP as control. We identified stable loop structures defining contact points between genomic loci pairs (loop anchors) from the contact frequency map (Hi-C map) (Figure S2C). Overexpression of the AS-RNA resulted in a decrease in total loops (Figure S2A). Lenti-GFP samples had an average detection of 86,382 significant interactions (loops), while Lenti-AS samples had 68,901 loops on average (Figure S2A). Loops in the AS-RNA samples made shorter range interactions compared to control samples (Figure S2B). Next, we performed differential loop analysis to identify changes in putative loop interactions that are due to the AS-RNA overexpression. These were labelled as static, gained and lost loops according to changes in interaction frequency. Though most loops/interacting loci were static, we found 545 loops that changed following AS-RNA overexpression (Figure 3A). 104 loops were gained and 441 lost. The dynamic loops made shorter range interactions than static loops (Figure 3B). Overall, most loops lost interactions, and this was consistent across all chromosomes (Figure S2D).

**Figure 3:**
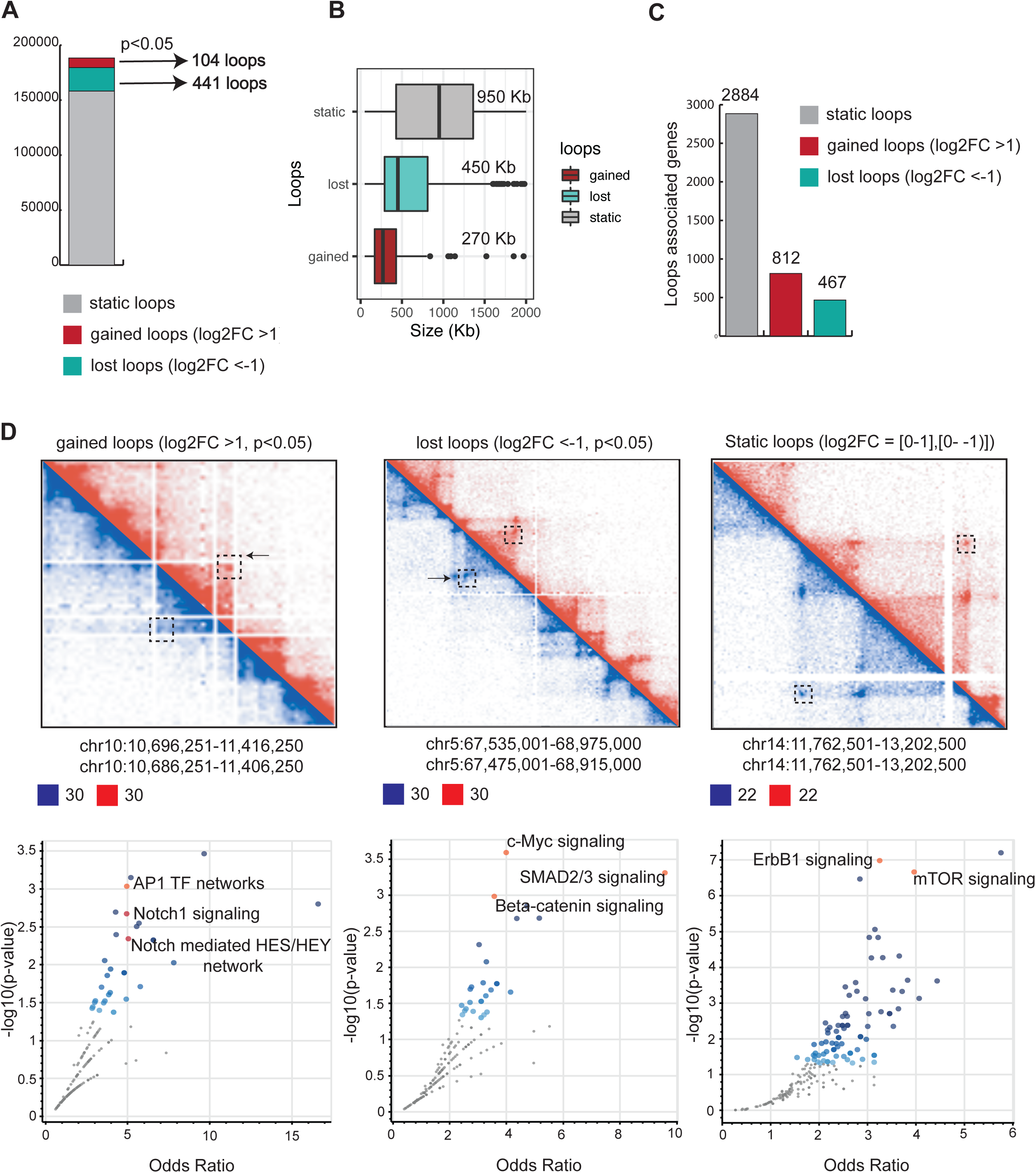
Hi-C following expression of the *EGR2*-AS-RNA in SCs shows 3D genome reorganization. A - C) Histograms showing the significant total loops called by the HiC+ software. Static loops (not gained or lost) are colored gray, lost loops are colored green and gained loops red (log2FC cutoff = 1, pvalue < 0.05). D) Detail of the Hi-C maps on the chromosomes 10, 5 and 14 showing examples of gained, lost and static loops respectively. The annotated loop anchors to the promoter of genes are represented by their main pathways (plotted as odds ratios by their corresponding p values).

Putative, stable loops generally represent sites of regulation and may facilitate cis-regulatory element to cis-regulatory element, promoter to promoter or cis-regulatory element to promoter interactions. We proceeded to annotate gene promoters existing at the loop anchors (Figure 3C). We found that promoters annotated to the gained loops were enriched for the AP-1 TF network and NOTCH1 signaling (Figure 3D). Lost loops were enriched for C-MYC signaling, β-CATENIN and SMAD2/3 signaling (Figure 3D). Finally, static loops were found to be enriched for ERBB signaling and mTOR signaling (Figure 3D).

### Integrated 3D genome reconstruction reveals clusters of cis-regulatory elements (CORES) and associated promoters

Hi-C defines loops that are 1) either structural in nature or 2) function to bring together DNA regulatory elements. To identify the latter, we found loops whose contact point (anchors) involves a promoter and its regulatory element (Figure 4A). Previous work has shown that DNA regulatory regions such as promoters, enhancers, silencers and insulators are in open chromatin, nucleosome depleted areas ^7^. A limited set of such accessible sites are composed of large intergenic clusters of accessible chromatin sites (<20 Kb apart) that are cell type specific, hence reminiscent of super enhancers ^8–10^.

**Figure 4:**
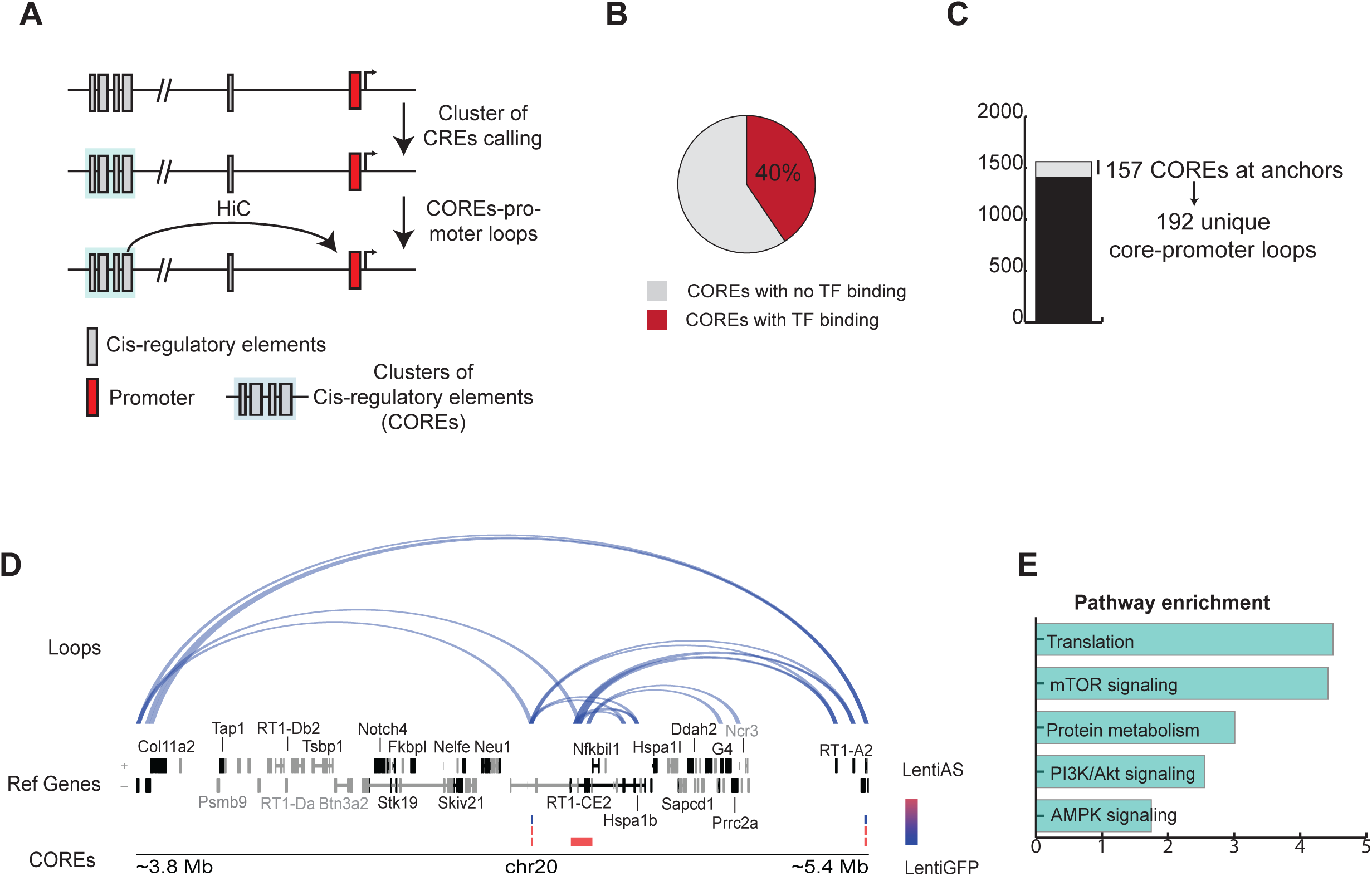
Reconstruction of long-range interactions between Cis-regulatory elements (COREs) and associated promoters. A) Schematic of clusters of regulatory elements (COREs) from chromatin accessibility data (ATAC-seq). Clusters of multiple proximal cis regulatory elements (CREs) were used to define COREs. Interacting COREs and promoters were annotated to loop anchors. B) Sub-setting of COREs based on transcription factor binding. 40% of COREs are occupied by at least 1 TF footprint. C) Hi-C reveals 192 COREs-promoter loops. 157 of 1563 COREs form long distance interaction with promoters. D) Example of COREs-promoter interactions at a genomic region at chromosome 20. COREs are colored by samples, red and blue for the AS-RNA and GFP control, respectively. Reference genome is color coded for expressed and non-expressed genes, respectively black and grey. E) Pathway enrichment analysis of COREs interacting genes.

These regions were dubbed “clusters of cis-regulatory elements” or COREs. Using an unsupervised machine learning approach, we identified an average of 309 and 472 of such clusters (CORES) in control and AS-RNA samples, respectively (Figure S3A). Additionally, we noted a heavy TF occupancy at 40% of the COREs with a median occupancy of 155 footprints (Figure 4B, Figure S3C). These data cast half of the COREs as being highly active regulatory DNA sites.

Distal DNA regulatory elements with significant TF occupancy are known to form regulatory, stable loops at promoters. To identify these loop structures, we identified loops whose anchors join a distal CORE and its target promoter(s). Approximately 10% (157/1563 total CORES across all samples) contacted at least one promoter site (Figure 4C & 4D). There were in total 192 long-distance genomic interactions between CORES and promoters (Figure 4C). These interactions occurred at median genomic distances of 365 Kb and 340 Kb in GFP control and AS-RNA respectively (Figure S3B). Pathway enrichment analysis of the CORES interacting genes showed top enrichment for translation, mTOR signaling, protein metabolism, PI3KC1/AKT signaling and AMPK signaling (Figure 4E).

### Expression of the *EGR2*-AS-RNA induces regulatory changes between the *mTOR* **promoter and its cis-regulatory elements**

Of the 192 COREs-promoter interactions, we focused on regulation at the *mTOR* promoter. Beyond its established ^11^ significance in SC plasticity, we elected to further probe regulation of the *mTOR* promoter since it interacts with a large cluster of regulatory elements (a region covering ∼100 Kb) reminiscent of a super enhancer (Figure 5A); and the *mTOR* region was one of few regions forming multiple contacts with a CORE (8 interactions total), potentially suggesting a regulatory hub (Figure 7A). We tested whether this regulatory region uniquely interacts with the *mTOR* region by screening for all other possible interactions and found none. This suggests that we discovered an *mTOR* regulating super enhancer/regulatory region located ∼336 Kb bases away from the *mTOR* promoter (Figure 5A).

**Figure 5:**
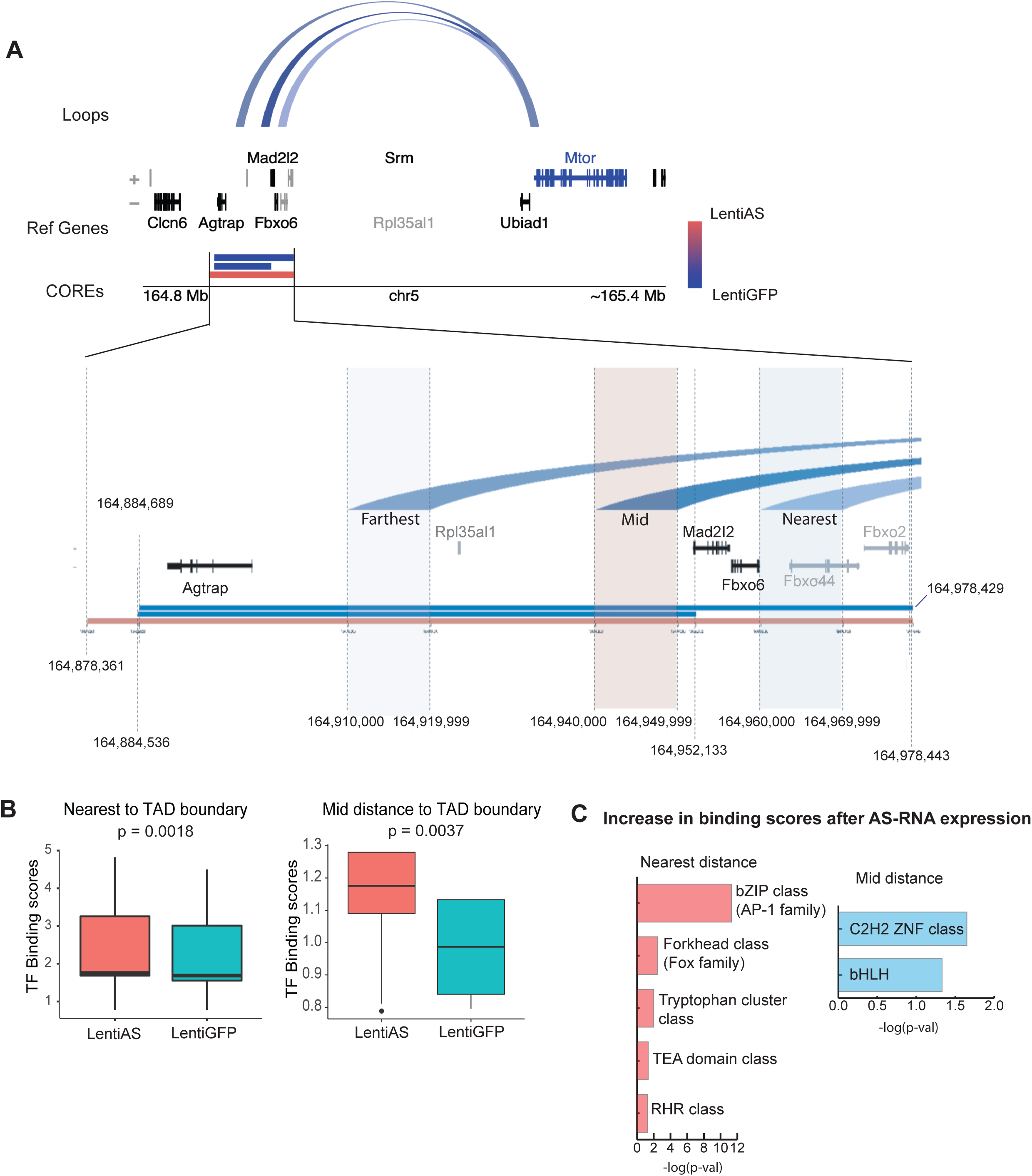
Expression of the *EGR2* promoter AS-RNA results in changes between the mTOR promoter and its cis-regulatory elements. A) Depicts the three contact points completely overlapping COREs and mTOR promoter. Magnification of the COREs region with anchors in blue with detailed coordinate characterization of the region. Characteristic of loop anchor region overlapping the COREs region. “Nearest/mid/farthest” distance to TAD boundary nomenclature denotes the three loop anchors position within the TAD in relation to the closest boundary. COREs are colored by samples, red and blue for AS-RNA and GFP control, respectively. Reference genome is color coded by expressed and non-expressed genes, respectively black and grey in color. mTOR shown in blue in gene reference panel. B) Change in TF binding scores of all TFs at nearest and mid distance to TAD boundary loop anchors following expression of the EGR2-AS-RNA (farthest to TAD boundary loop anchor could not be analyzed due to very low occupancy). C) TF families that experience the greatest increase in binding scores following expression of the AS-RNA.

Lower resolution inspection of contact frequencies at the COREs-*mTOR* locus (500 Kb-∼1 Mb resolution) revealed that *mTOR* and its regulating super enhancer region are in different domains (Figure S4A, Figure S4C). Closer inspection of COREs-*mTOR* promoter interaction showed three main points of contact that originate from the COREs region and perfectly contact the *mTOR* promoter (Figure S4A, Figure 5A). We further differentiate these contact points based on distance from the nearest TAD boundary, labelled “nearest distance”, “mid distance” and “farthest distance” to TAD boundary (Figure 5A, Figure S4B). These contact points harbored significant TF binding with 408 footprints on aggregate (Figure S4B). We noted increasing TF occupancy as the contact sites approached the TAD boundary (Figure S4B). Next, we clustered the TFs present at the “nearest distance”, “mid distance” and “farthest distance” to TAD boundary in TF communities based on the presence of families of TFs. The farthest distance to boundary interaction showed only REST occupancy, a well-known repressive transcription factor in neuronal lineage cells (Figure 6C). The mid distance to TAD boundary had 65 TFs with an enrichment for the C2H2 zinc finger class of TFs, such as GLI3, GLI2, GLIS3, GLIS2 and MAZ (Figure 6B). The nearest to TAD contact had 342 TF with a predominance for the AP-1 family of TFs, structural TFs such as CTCF, VEZF1 and MAZ and for zinc fingers of the Kruppel family (Figure 6A). Finally, we found a significant increase in TF binding at the nearest and mid-distance to TAD boundary sites following AS-RNA expression (Figure 5B). Specifically, expression of the AS-RNA induces significance increase in TF binding of the AP-1 family, FOX family, Tryptophan cluster class, TEA domain class, RHR class nearest to the TAD boundary sites and significant increase in binding of the GLI family and C2H2 TFs at the mid-distance to TAD boundary sites (Figure 5C).

**Figure 6:**
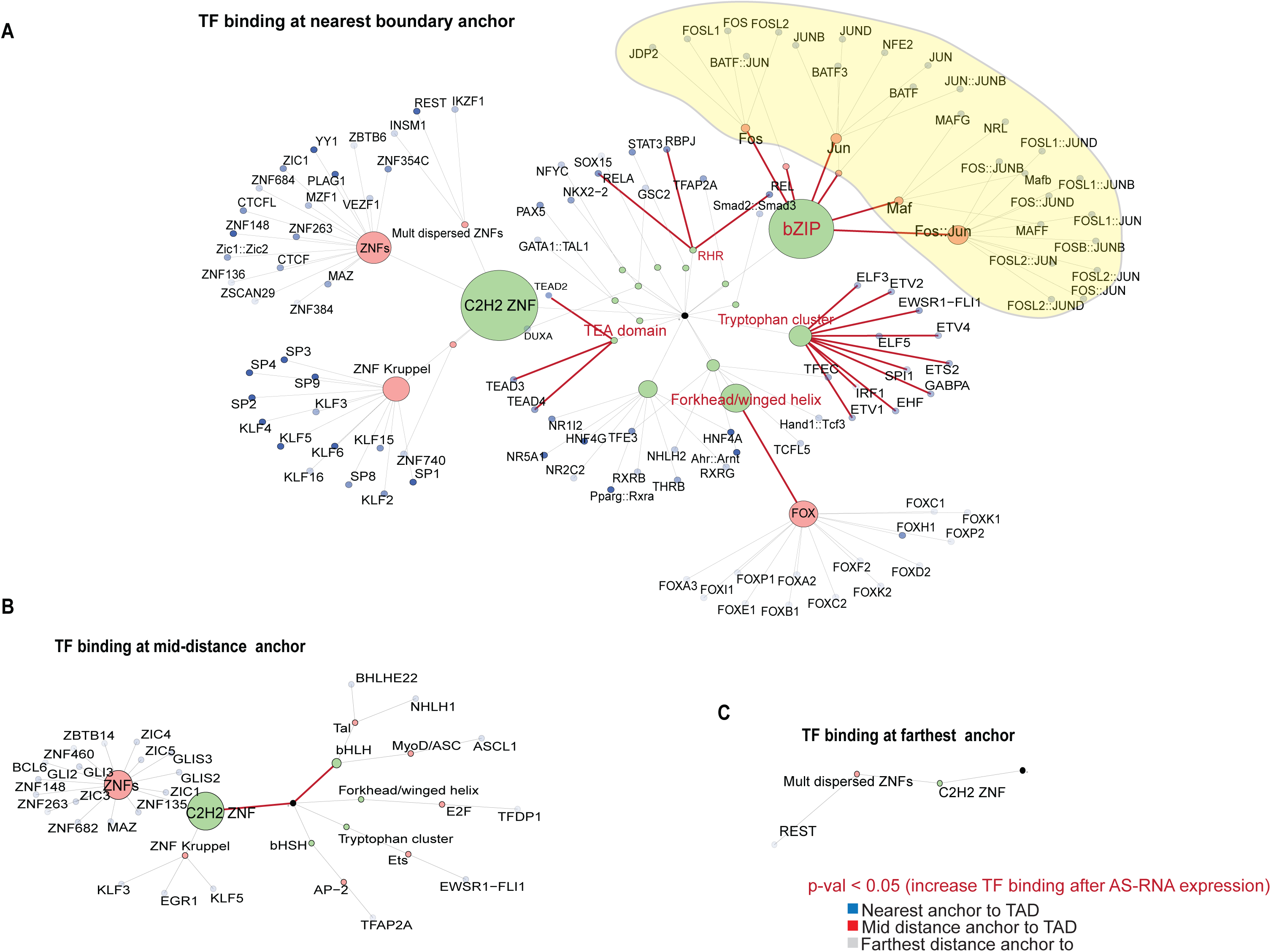
Network of TF binding dynamics at the mTOR interacting regulatory element/super enhancer. A) Network of TF binding at the loop interacting with the COREs at the nearest TAD boundary. B) Network of TF binding at the loop interacting with the COREs at the mid distance to the TAD boundary. C) Network of TF binding at the loop interacting with the COREs farthest to the TAD boundary. Maroon color emphasizes TF class and families that experience a statistically significant increase in TF binding following expression of the *EGR2*-AS-RNA (Wilcoxon test, p-val <0.05).

### The *EGR2*-AS-RNA induces changes at the *mTOR* interdomain regulatory hub

We determined that the COREs-*mTOR* interactions occur via an interdomain contact (Figure S5A). While most of all interactions occur within TADs, 30% of interactions occur between TADs (Figure S5D). Additionally, we found that these interdomain interactions occurred mostly in non-adjacent TADs (Figure S5C and S5D). The *mTOR* and COREs harboring TADs, which both occupy active compartments (A compartments), exhibit preferential interdomain interactions (Figure S5A, Figure S5B). We detected 8 total interactions linking the CORE containing TAD and the *mTOR* containing TAD (Figure 7A). This was one of the highest noted interactions between a CORE and a gene region. Additionally, we noted >400 footprints at *mTOR* promoter contacting COREs, far superseding the median occupancy of 155 footprints found at all COREs across all samples (Figure S3C). The stability of the interactions and the high TF occupancy suggested a heavily regulated hub between the *mTOR* promoter and its interacting super-enhancer region.

**Figure 7:**
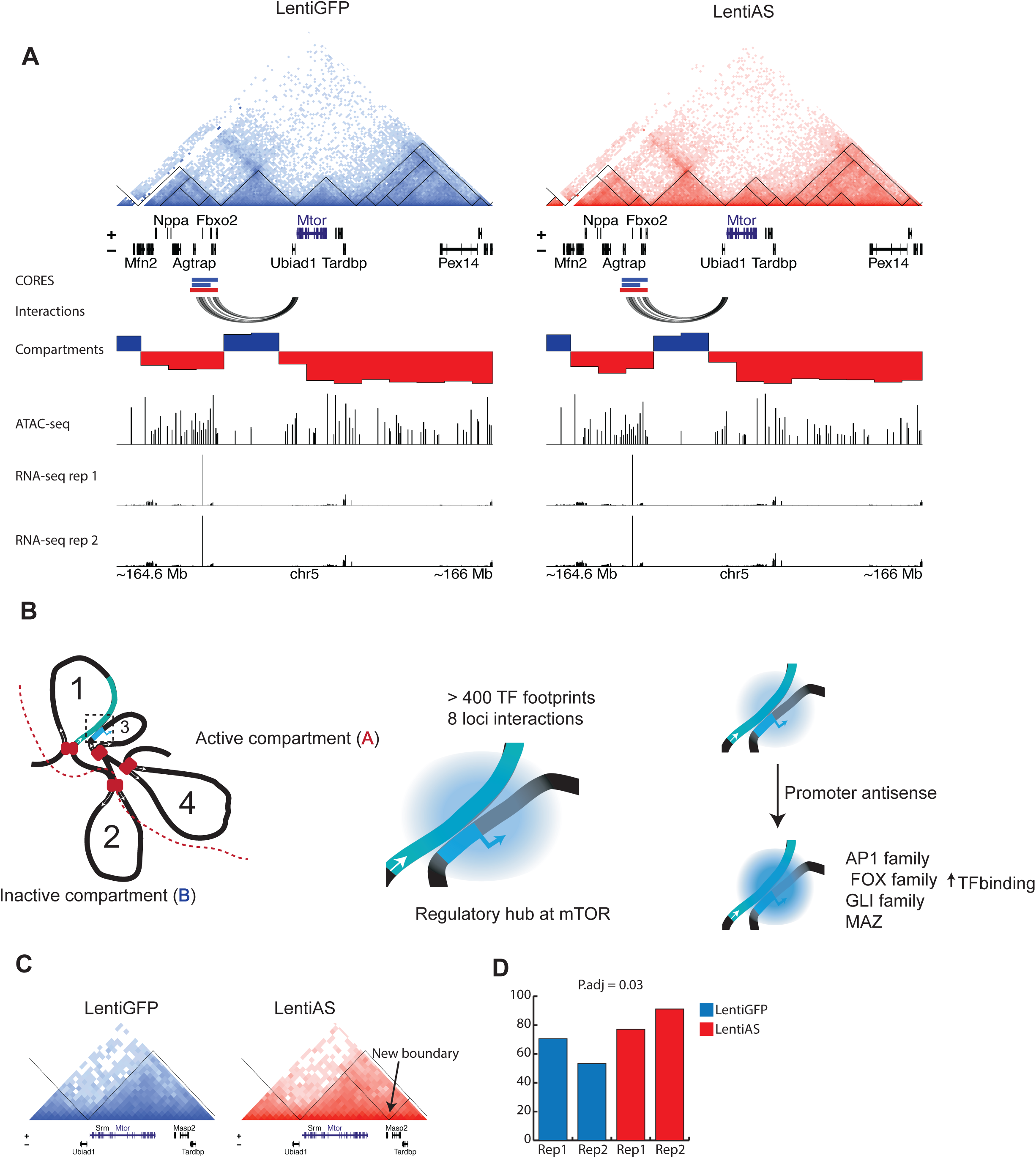
*EGR2*-AS-RNA induces structural reorganization and TF binding changes at the mTOR interdomain regulatory hub. A) Depiction of mTOR interaction hub harboring meta-TAD. Tracks shown below (from top to bottom): clusters of cis-regulatory elements (COREs), genomic loops, active/inactive compartments, ATAC-seq reads, RNA-seq reads for individual replicates B) Proposed model of 2D interdomain interactions between COREs and mTOR promoter. Magnified depiction of activity at hub shown with blue intensity. C) Depiction of changes at mTOR harboring TAD. D) Bar-plots of normalized gene expression in control and AS-RNA expressing cells (p.adj = 0.03).

Next, we probed changes occurring at this *mTOR* regulatory hub following AS-RNA expression. First, there was an overall increase in TF binding at the hub following AS-RNA expression, with significant increase in binding scores of the AP-1 TF family, FOX, GLI and MAZ TFs (Figure 7B).

Second, we noted the formation of a new boundary at the 3’ end of the *mTOR* gene after AS-RNA expression (Figure 7C). This new boundary resulted in the formation of a nested sub structure, whereby *mTOR* is more insulated from other possible intra and inter TAD interactions (Figure 7C). Such changes may function to maximize *mTOR*-CORE interaction, while minimizing other interactions. The TF activity and structural changes at the *mTOR* interaction hub were accompanied with moderate, but statistically significant increase in *mTOR* expression levels following AS-RNA expression (Figure 7D).

## Discussion

Recently, we demonstrated that an *EGR2* promoter antisense RNA is induced following in vivo sciatic nerve injury. The AS-RNA inhibits *EGR2* transcription via recruitment of components of the PRC2 complex (EZH2, H3K27me3) to the *EGR2* promoter, while inhibition of the AS-RNA expression delays demyelination in a sciatic nerve transection model ^5^. It has been shown that long non-coding RNAs can have opposing regulatory effects by simultaneously binding to epigenetic activators and inhibitors ^12, 13^ but this role has not been established for promoter antisense RNAs. We show here that the *EGR2*-AS-RNA recruits activating and repressing histone marks on the promoters of *C-JUN* and *EGR2* respectively and increases chromatin accessibility in promoters that exhibit increased binding of AP-1 family TFs. Since expression of the EGR2 promoter AS-RNA induces chromatin remodeling, we questioned the possibility that expression of the AS-RNA could affect spatial genome organization, which could result in global transcriptional regulation from a single epigenetic input.

Recently it was shown that subtle changes in chromatin loop contact can lead to significant changes in gene expression ^14^. These loops can bring together the regulatory elements with their target sequences that can be distant from one another. After the AS-RNA expression, we observe a decrease in the total number of loops formed, and these loops make shorter range interactions relative to control samples. It has been suggested that larger loops may have a structural role and could therefore be involved in the formation of insulated neighborhoods to protect and stabilize interactions between promoters and regulatory elements, while shorter range interaction loops may be more regulatory ^15–17^. In addition, our Hi-C data show enrichment of genes involved in AP-1 and NOTCH1 signaling at the gained loop anchors suggesting that AP-1 and NOTCH1 signaling network experience a change in regulation due to increased contact with distal regulatory elements following expression of the AS-RNA.

Clusters of cis-regulatory elements (COREs) are reminiscent of super-enhancers, which are large clusters of active enhancers that drive gene expression and confer specific cell identity ^18^. We identified COREs with high TF activity, suggesting that these are hubs for transcriptional regulation. These CORES form long range interactions with promoters of genes that belong to the mTOR, AKT, AMPK and protein translation pathways. Areas of the genome where chromatin interactions are more frequent are termed “topologically associating domains” (TADs) ^19, 20^. The positions of TADs within the genome are stable between several cell types and even across species ^21, 22^ suggesting that TADs are architectural units that house regulatory interactions. Recent work has shown the existence of a certain hierarchy with TADs where domains are included within other domains (meta-TADs) through TAD-TAD interactions ^23, 24^ and this level of organization correlates with cell specific transcriptional and epigenetic regulation. However, the molecular mechanisms that regulate this cell-specific architecture are not known. In addition, the possibility that changes in TAD-TAD interaction frequencies could have profound effects on cell identity programs open exciting possibilities for understanding and even modifying cell specific responses. To gain insight into the potential role of the *EGR2* promoter AS-RNA in modulating interactions between different TADs, we focused on the CORE-promoter interaction of *mTOR* since this CORE has all attributes of a regulatory hub.

Low resolution analysis of our Hi-C data following expression of the AS-RNA shows that the promoter of *mTOR* and its regulatory COREs are positioned in separate TADs and form three distinct loop contacts. Expression of the AS-RNA significantly increases TF binding of AP-1, FOX, Tryptophan class, RHR class and GLI families that form TF communities on the inter-TAD contact boundaries between *mTOR* promoter and its COREs. In addition, expression of the AS-RNA induces formation of a new boundary at the *mTOR* gene that insulates *mTOR* from other intra or inter TAD interactions. The role of *mTOR* varies on SCs depending on the cellular state: it is crucial for myelination/differentiation of SCs during development ^25, 26^, but in adulthood mTOR has been proposed to contribute to SCs remarkable plasticity ^11^. It has also been described as one of the first proteins that are upregulated after nerve injury, promoting C-JUN elevation and SC dedifferentiation ^27^.

We propose here that promoter AS-RNAs could function as molecular recruiters of histone modifying enzymes and may regulate interdomain regulatory hubs that are cell specific. Our results raise some intriguing possibilities and interesting mechanistic questions. Are these meta-TAD interactions formed by chromatin folding on single cells or are they results of aggregate populations of cells? Single-allele chromatin interactions do reveal regulatory hubs ^28^, supporting the notion that these interactions occur in individual cells. What directs the binding of AS-RNA-histone mark complexes on specific TADs housing regulatory hubs? The presence of large communities of TFs suggests that there is crosstalk between TF regulation and promoter AS-RNA production. The possibility of TF feedback regulation of the promoter AS-RNA should be explored, and it could explain cell specific autonomous regulation. Finally, the ability of the *EGR2* promoter AS-RNA to regulate spatial genome architecture in SCs opens exciting possibilities for development of RNA therapeutics targeting promoter AS-RNAs for nerve regeneration.

## Acknowledgements

This work was supported by a Warren Alpert Foundation grant (Project #17775) and internal funds of the Neurosurgery Department to N.T. Part of this work was facilitated by the Brown University Genomics Core.

## Materials and Methods

**Methods for cell cultures, Transfections, qPCR, TF activity assays and Primer sequences are included in the Supplemental Information.**

### RNA-seq

Sequencing reads were aligned to the rn6 genome assembly with hisat2 (Should this be hisat2 instead of hista2?) ^29^. Gene-wise read summarization was done with featureCounts using Refseq exon coordinates as specified in the rn6 refGene file available in the UCSC Genome Browser database ^30, 31^. Differential gene expression analysis was done in R via DEBrowser the DESeq2 platform ^32, 33^. We used an adjusted p-value cut-off of 0.05 and a fold change of 1.3.

### ATAC-seq

Sequencing reads were aligned to the rn6 genome assembly with bowtie2, and duplicate reads were removed with picard-tools ^34, 35^. Promoters were defined as 2000 bp regions positioned between −1500 to +500 nucleotides with respect to the transcription start site, as annotated in Refseq. Peaks of chromatin accessibility were detected using MACS2, and those located at promoter regions were identified using the intersectBed command in BEDtools (previously “intersectBed routine of Bedtools”). ^36, 37^. To compare relative promoter accessibility, ATAC reads mapping promoters were quantified with featureCounts and the promoter locations defined above. Differential promoter accessibility was determined with the limma package ^38^. A p-adjusted value of 0.05 and fold change of 1.5 were used as cut-offs.

### RNA immunoprecipitation (RIP)

To perform RIP, we used the magnetic RNA ChIP kit. Briefy, cells were subjected to crosslinking in 1% formaldehyde in PBS, then lysed in the kit-provided lysis buffer, and then, after centrifugation, the pellet that contained the chromatin was sheared to 100-1000 bp (the settings of the sonicator depend on the cell type or tissue type). Sheared chromatin was subjected to DNAse treatment and then we proceeded to the immunoprecipitation with the magnetic beads. For each immunoprecipitation, we added 10 µg of sheared chromatin and 1 µg of ChIP grade antibody (list of antibodies in table 2) and an isotype control for unspecific binding to the magnetic beads. Then we incubated the tubes to an end-to-end rotator overnight at 4°C. Then, we eluted the RNA and proteins, degraded the proteins with Proteinase K and reversed the crosslinks. We purified the RNA doing a Trizol extraction and performed qPCR against AS-Egr2 RNA. Relative enrichment for each experimental sample was calculated as a percentage of the input, and the isotype control percentage was deducted from each experimental condition.

### Chromatin Immunoprecipitation (ChIP)

For DNA Chromatin Immunoprecipitation (ChIP) we used the ChIP-IT High Sensitivity^®^ kit from Active motif. Lysates were obtained as described in the RIP protocol. Lysates were incubated overnight at 4°C on rotation with the specific ChIP-verified antibodies (list on table 2). For negative control ChIP, we used a non-targeting isotype-matched IgG. Chromatin was then precipitated, and DNA was extracted. Recovered material from the input sample and all the ChIP samples per condition were used to perform qPCR of the Egr2-AS-RNA. Calculations were performed as the RIP experiments.

### Hi-C capture method

Hi-C was performed using the Arima-HiC kit. As per their protocol, we tested the amount of input sample beforehand and we used 2 million cells per each independent sample. We did QC on samples in every recommended step. Sonication was performed using the Covaris s220 instrument to a fragment size of 300-400 bp (Settings: Power: 5, Duty Factor: 10%, Cycles per Burst: 200, and Time: 60 sec per process). We used KAPA Hyper Prep Kit to generate libraries. The primers used for indexing were obtained from Illumina, and the libraries were quantified and QC was done using the KAPA Library Quantification Kit. Libraries were sequenced by GENEWIZ using the Illumina HiSeq 2500 system to acquire 150 bp paired-end sequence reads, reaching 300M reads for each sample based on total usable reads >20 Kb.

Raw and ICE-normalized contact matrices were prepared with HiC-Pro (Servant et al., 2015). ICE-normalized and condition-merged HiC-Pro matrices were converted to Juicer.hic files for resolutions 10 Kb and 100 Kb using a wrapper for the Juicer tools Pre command in the HiC-DC+ Bioconductor package ^39, 40^.

### Significant and differential loop analysis

Significant loops (q-value <= 0.05) and differential loops (log2FC cut-off = 1, p-value < 0.05) across conditions were identified using HiC-DC+ with raw HiC-Pro matrices and bed files of two AS replicates and two GFP replicates at 10 Kb resolution ^40^. To filter biases in denser genomic regions in close proximity and noise in sparser regions that are more far apart, only loops between 50 Kb and 2 Mb were called.

We categorized the loops into three sets: 1) static for significant loops that did not meet the differential loops thresholds, 2) dynamic gain for significant loops that were differential with a log2FC > 1, and 3) dynamic lost for significant loops that were differential with a log2FC < −1. Dynamic loops represent interactions that, after antisense overexpression, were gained or lost relative to their interaction frequencies in GFP control samples.

### Identification of clusters of cis-regulatory elements (COREs)

The R package CREAM version 1.1.1 was used to identify clusters of *cis*-regulatory elements, or COREs ^8^. The bed files for the peaks were used as an input and the program was run with parameters MinLength = 1000 and peakNumMin = 2 for all the LentiAS and LentiGFP samples.

### Annotation/identification of COREs-promoter loops

Gene symbols obtained from the RNA-seq analysis were first mapped to stable rn6 Entrez Gene identifiers, then to transcripts, and finally to 2200 bp-width promoters using the Bioconductor packages TxDb.Rnorvegicus.UCSC.rn6.refGene, BSgenome.Rnorvegicus.UCSC.rn6, and org.Rn.eg.db ^41–43^. We defined COREs- promoter loops as loops that consisted of one anchor intersecting with at least one promoter and the other anchor intersecting with at least one COREs. Anchors of COREs- promoter loops had at least a 1 bp-overlap with promoters or COREs. A subset of these loops had complete overlaps at anchors in which an entire promoter or COREs was contained in the 10 Kb loop anchor or, for COREs greater than 10 Kb in width, the entire loop anchor was contained in the COREs.

### Genome wide differential TF activity at promoter sites

Peaks were annotated against the rn6 genome. The R package diffbind from Bioconductor was used to compare the similarity in peak read counts via correlation heatmaps and principal components analysis ^44^. The R package DESeq2 from Bioconductor was used to detect differential peak regions ^33^. To do so, a consensus set of peaks was first generated via the runConsensusRegions function in the *soGGi* package (Dharmalingam and Carroll, 2015), and then read counts of each peak range in the consensus peak set were calculated via the featureCounts function in the Rsubread package for each sample ^31, 45, 46^. The resulting counts matrix was then processed via DESeq, and differentially accessible chromatin regions between control and experimental samples were defined as peak regions with an absolute fold change greater than 1.5 and a p-value less than 0.05. Genes associated with differentially accessible peaks were identified through a many-to-many mapping algorithm implemented in the seq2gene function from the ChIPseeker package ^47^. Promoters with differential accessibility were defined as differentially accessible peak regions within 3000 bp upstream and downstream of a TSS in the rn6 genome. DAStk was used to quantify significant differences in transcription factor activity at the differentially accessible promoter sites ^48^. Motif binding sites with differential activity were analyzed through EnrichR to identify enriched gene ontology categories ^49^.

### Site-specific TF footprint/binding predictions

TOBIAS was used to identify specific footprint location at DNA regulatory elements ^50^. First, the ATACorrect function was ran to correct for tn5 insertion bias. Next, depletion signal (negative counts) and general accessibility at surrounding regions were used to derive a binding score. Finally, to make TF binding activity predictions, the footprint scores were matched to binding motifs from the 2020 JASPAR CORE database ^51^. TF binding activity were restricted to accessible loop anchor sites using genomic contact regions derived from Hi-C data. Finally, the exact sites predicted to be bound were extracted for further site-specific downstream analysis.

### Site-specific TF binding predictions at regulatory elements

All intersections of TF footprints with anchors, promoters, and COREs had an overlap of at least 1 bp. We identified TF footprints that intersected with COREs and with all anchors overlapping with COREs. TF footprints that intersected with each of the three loop anchors interacting with the mTOR promoter (i.e. anchors containing COREs) were grouped into families and classes based on the 2020 and 2022 JASPAR CORE databases ^51, 52^. To identify significant differences between AS and GFP footprint scores (binding scores) at each anchor, we performed Wilcoxon rank sum tests in R on 1) the full list of TF footprints that intersected with the loop anchor and were present in both AS and GFP samples and 2) the same list grouped by TF class.

### Identification of TADs

The C++ software OnTAD was to detect hierarchical TADs in ICE normalized and condition-merged .hic files at 10 Kb resolution ^23^. The OnTAD executable file was compiled under gcc 8.3 and run with default parameters.

### Intra and inter TAD loops localization

We defined intra-TAD loops as loops with both anchors intersecting with the same outermost TAD and inter-TAD loops as loops with anchors intersecting with different outermost TADs. All intra- and inter-TAD loop anchors were annotated with genes with promoters that overlapped with any region of the anchor.

### Identification of A/B Compartments

The eigenvector command in Juicer tools version 1.19.02 was used to call A/B compartments in ICE-normalized and condition-merged .hic files at 100 Kb resolution ^39^. The signs of the output eigenvectors indicate the compartment, but the signs are arbitrary in each set of eigenvectors calculated for each chromosome and condition. Therefore, for regions of interest, eigenvectors were plotted alongside ATAC-seq narrow peaks using the plotgardener R package to determine whether they corresponded to compartment A or B ^53^.

### Software & Data availability

Hi-C analysis scripts are available at https://github.com/hyeyeon-hwang/HiC-Analysis-of-Egr2-AS-RNA.

All datasets (RNA-seq, ATAC-seq, Hi-C) have been deposited to the GEO Omnibus and can be accessed through: https://www.ncbi.nlm.nih.gov/geo/query/acc.cgi?acc=GSE201627

### Plasmid Availability

The lentivirus construct (LentiAS) is deposited to Addgene and is available to order (#177737).

## Supplemental Information

### Supplemental Experimental Procedures

#### Rat Schwann cell culture

Rat Schwann cells were cultured as described earlier (Martinez-Moreno *et al*., 2017). Briefly, cells were cultured in DMEM supplemented with 10% FBS, 4 μM forskolin, 5 ng/ml heregulin-β1 and antibiotics, in surface modified 75cc *Primaria* flasks. Cells were fed every other day and passaged at 80% confluence.

#### Transfection experiments

Transfection of RSC with the target vectors/siRNAs or the negative controls were performed with the TransIT-X2 transfection reagent. Briefly, we plated 2×10^5^ cells per well in a 6-well plate dish (*Primaria*, Corning). Then we directly applied the TransIT-X2 in Opti-MEM containing 1 microgram of each vector/25nM of each siRNA.

Overexpression of the Egr2-AS-RNA (and the backbone virus control carrying GFP) was performed with a lentiviral construct at a concentration of 2UFC/cell, with polybrene at a 1:1500 concentration, for 48 hours. GapMer against the Egr2-AS-RNA were added after 24 hours. The virus and GapMer sequences and generation were described in detail in previous work (Martinez-Moreno et al., 2017). GapMer sequence is proprietary of Exiqon (currently Qiagen). The plasmid to generate the lentivirus was deposited to Addgene and is available to order (#177737).

#### RT-PCR and qPCR

RNA was extracted from the cells, the sciatic nerves or the RIP eluates using Trizol and the PureLink RNA Mini Kit according to the manufacturer’s protocol, following a DNase treatment. 300 ng of RNA was reversed-transcribed to cDNA using SuperScript III First-Strand Synthesis System. For all qPCRs reported in the paper, we performed a no-reverse transcription (RT) control amplification to verify the absence of genomic DNA contamination with GAPDH primers. For the qPCR, we obtained the cDNA as explained above, and we ran a qPCR using SYBR Green PCR Master Mix with the same PCR primers at a final concentration of 250 nM. Relative mRNA levels were normalized to GADPH and quantified using the comparative Ct method, and fold change was calculated compared to each control. All primer sequences are shown in Table 1.

#### c-Jun transcriptional activity TransAM AP-1 c-Jun

We used the Nuclear Extract Kit and the TransAM AP-1 c-Jun kit (Active Motif). Briefly, we overexpressed the Egr2-AS-RNA in RSCs as explained above and we used 20 μg of the nuclear extract to run the transcription factor assay using the manufacturer’s recommendations.

### Supplemental Figure legends

**Figure S1:**
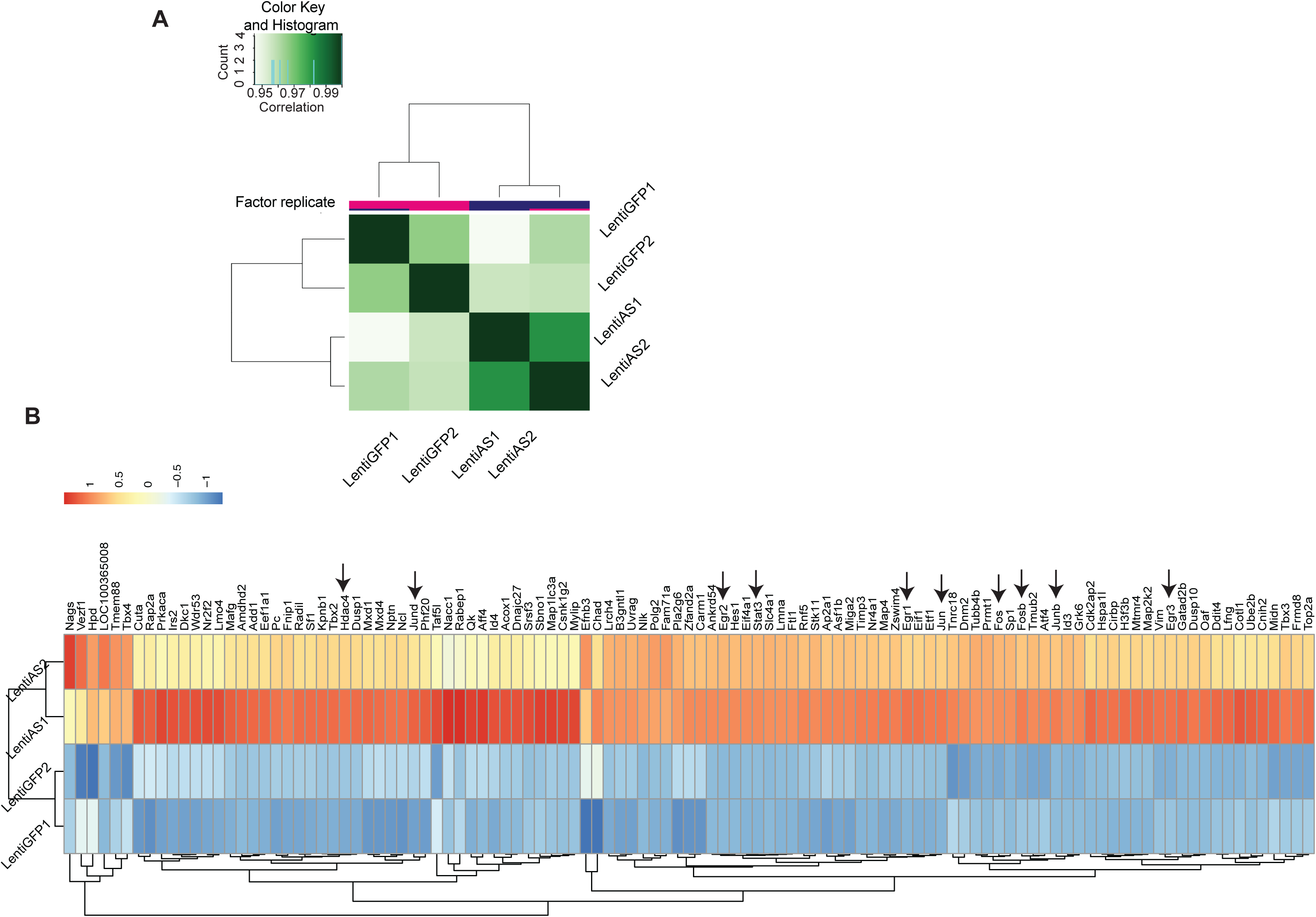
Expression of the *EGR2-*AS-RNA induces changes in chromatin accessibility. A) Similarity matrix analysis is shown as a heat map and the intensity of the color indicates cross-correlation between the compared groups. B) Heatmap of differentially accessible genes (defined as fold change > 1.5, with a P-value < 0.05).

**Figure S2:**
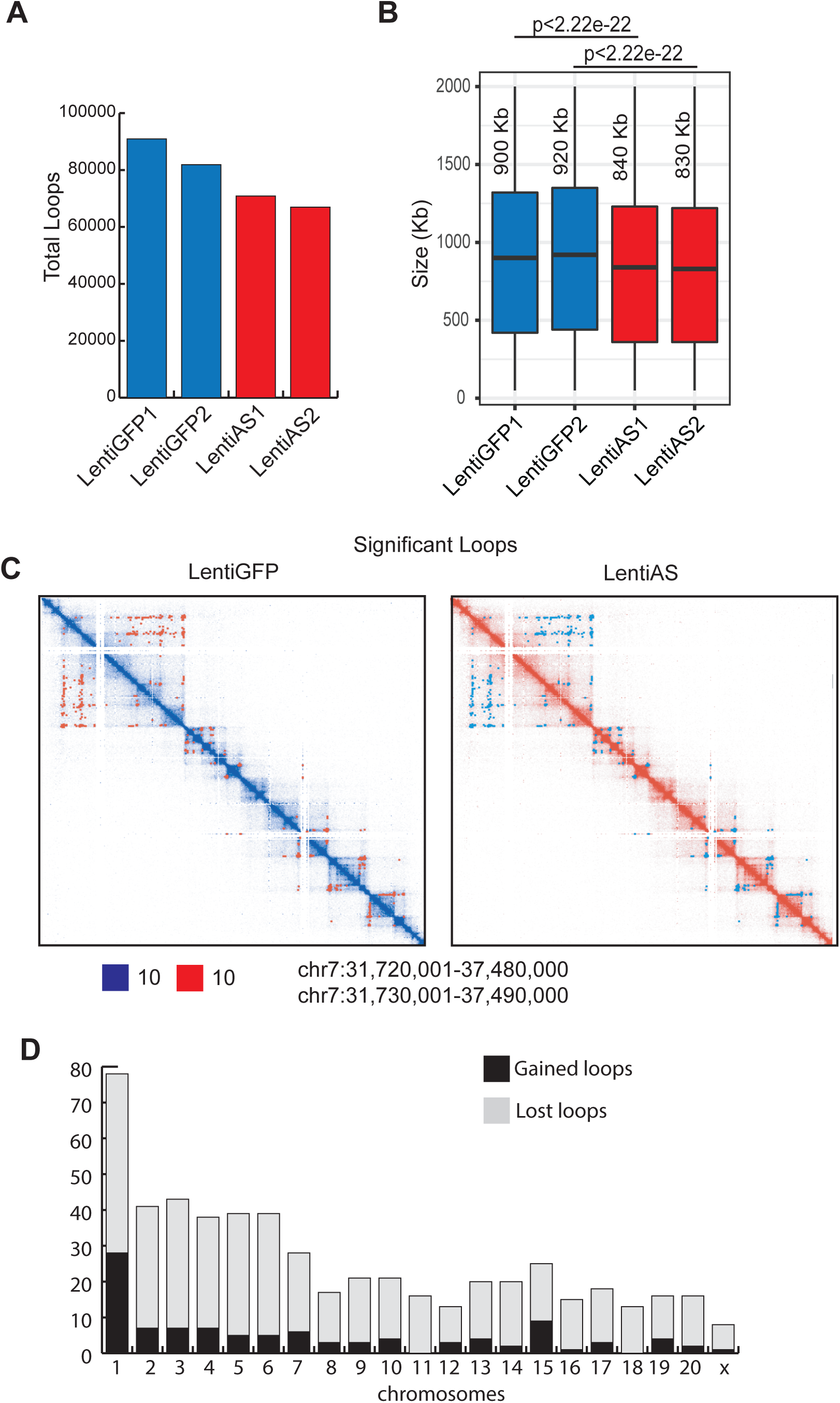
Loops description. A) Total number of loops per sample. B) Size of loops per sample. C) An example of significant loops detected in each condition. D) Gained and lost loops per chromosome.

**Figure S3:**
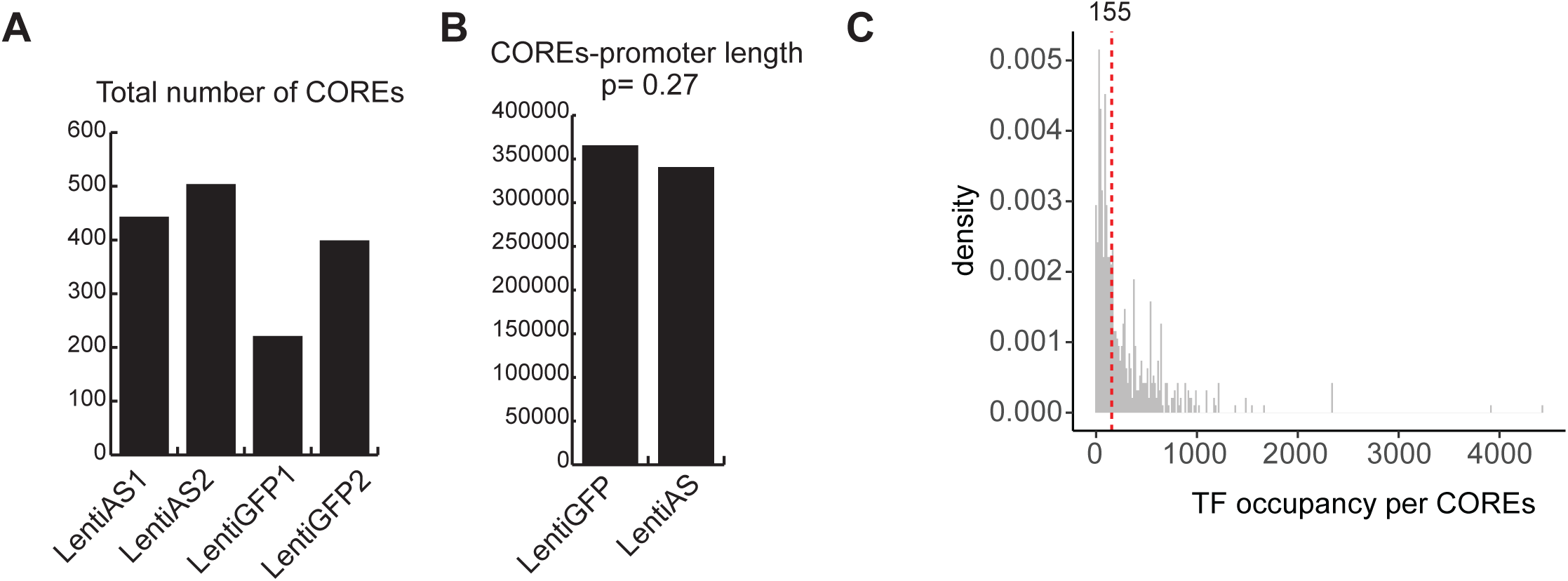
Characterization of COREs. A) Shows total number of COREs. B) COREs-promoter length between the AS-RNA expressing cells and GFP controls. C) Density plot of TF occupancy of all COREs across all samples.

**Figure S4:**
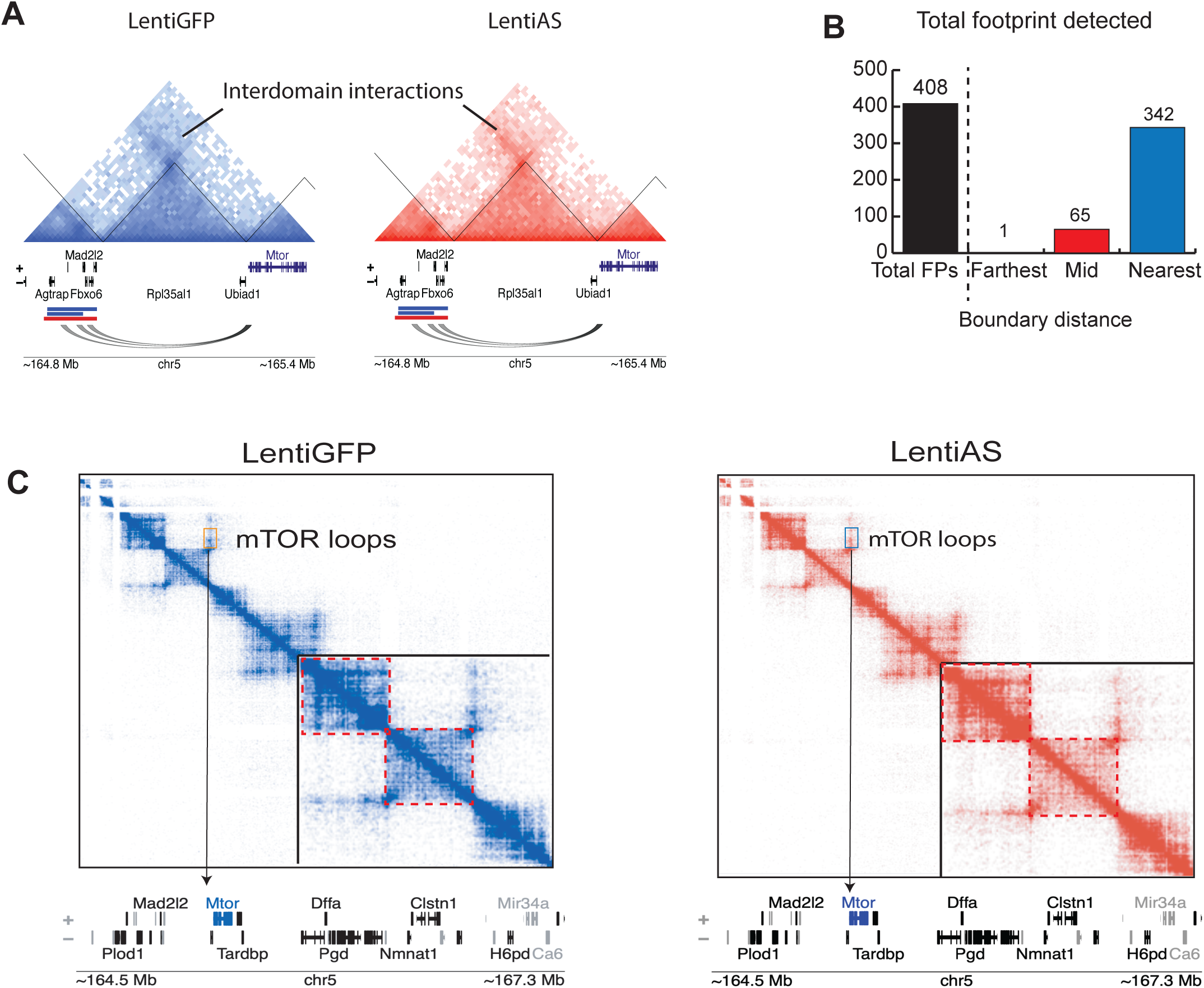
Characterization of loop interaction between mTOR and COREs. A) Depiction of TADs harboring mTOR and COREs in the AS-RNA and control samples. Interdomain interaction shown here as area of increased interaction intensity relative to surrounding bins. B) Total footprint detected at all loop anchors and shown by distinct loop anchors. Loop anchors named based on distance from nearest TAD boundary. C) Hi-C interaction map emphasizing interdomain mTOR loop interactions.

**Figure S5:**
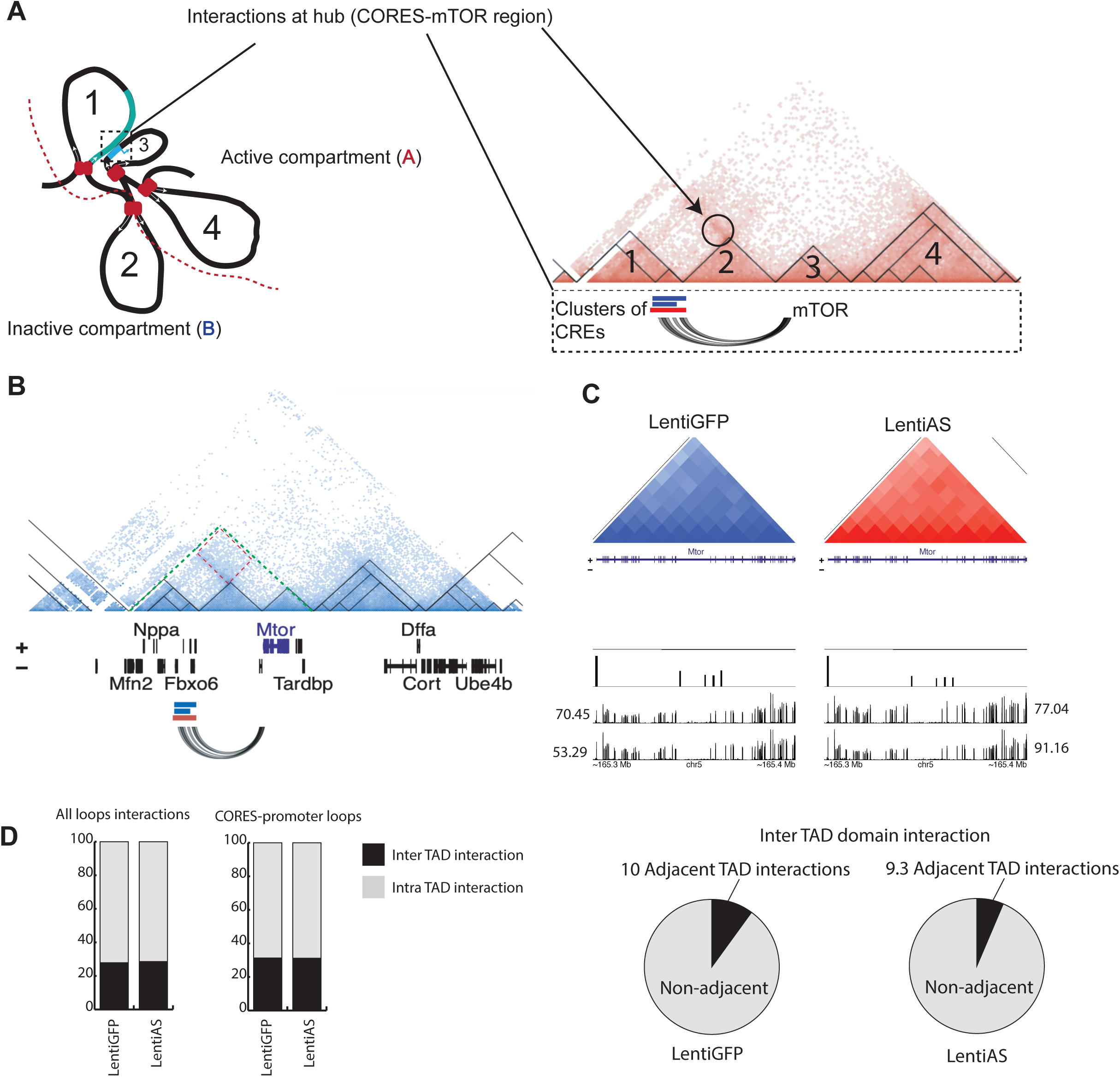
Characterization of mTOR interaction hub. A) Relates TAD interacting model to Hi-C data. B) MetaTAD region is emphasized with a green dotted line, while red box shows interdomain interaction. C) RNA-seq signal with normalized reads at the mTOR harboring TAD. D) Ratio of inter and intra TAD interactions between AS-RNA expressing cells and controls.

**Supplemental Table 1.**
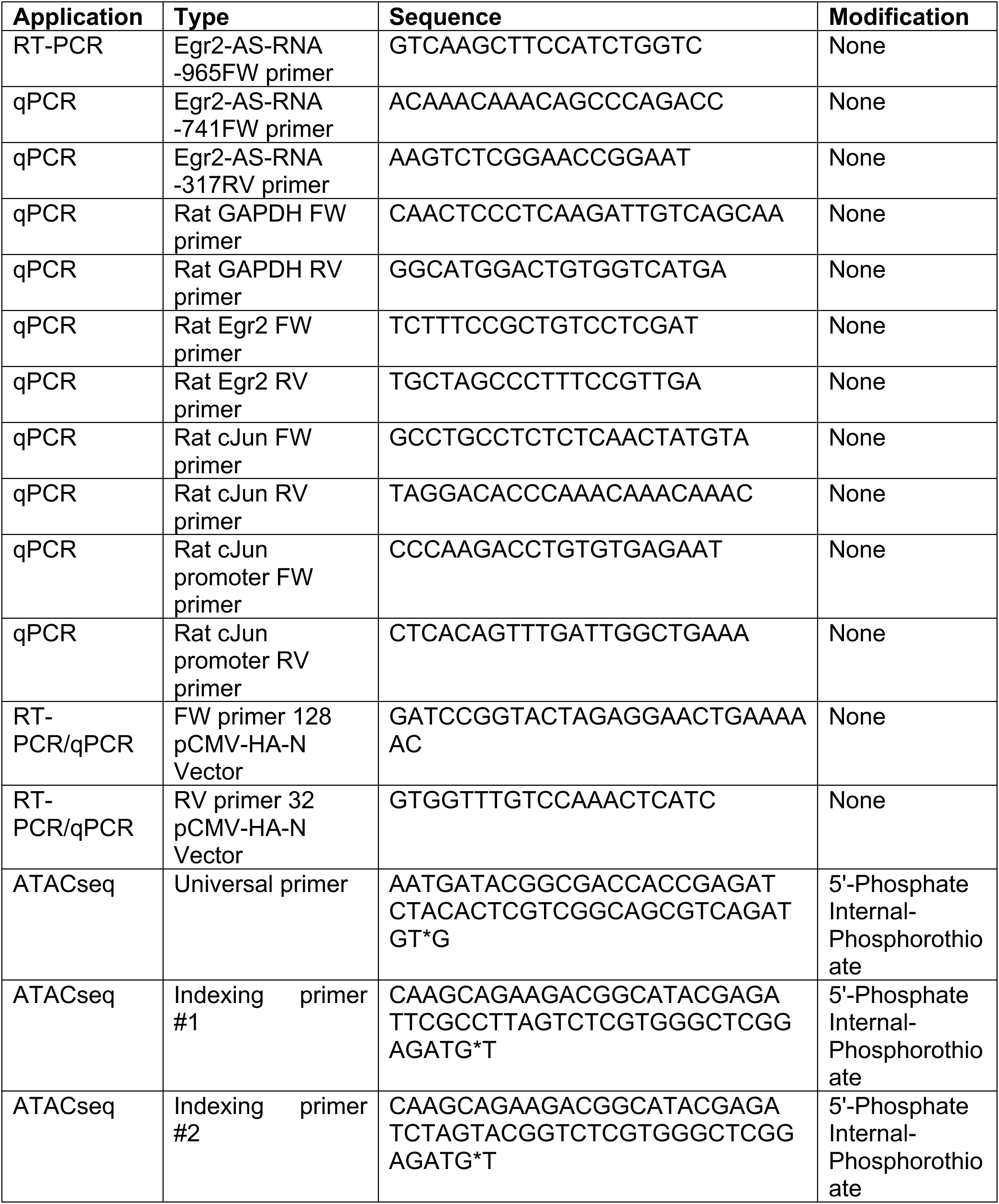
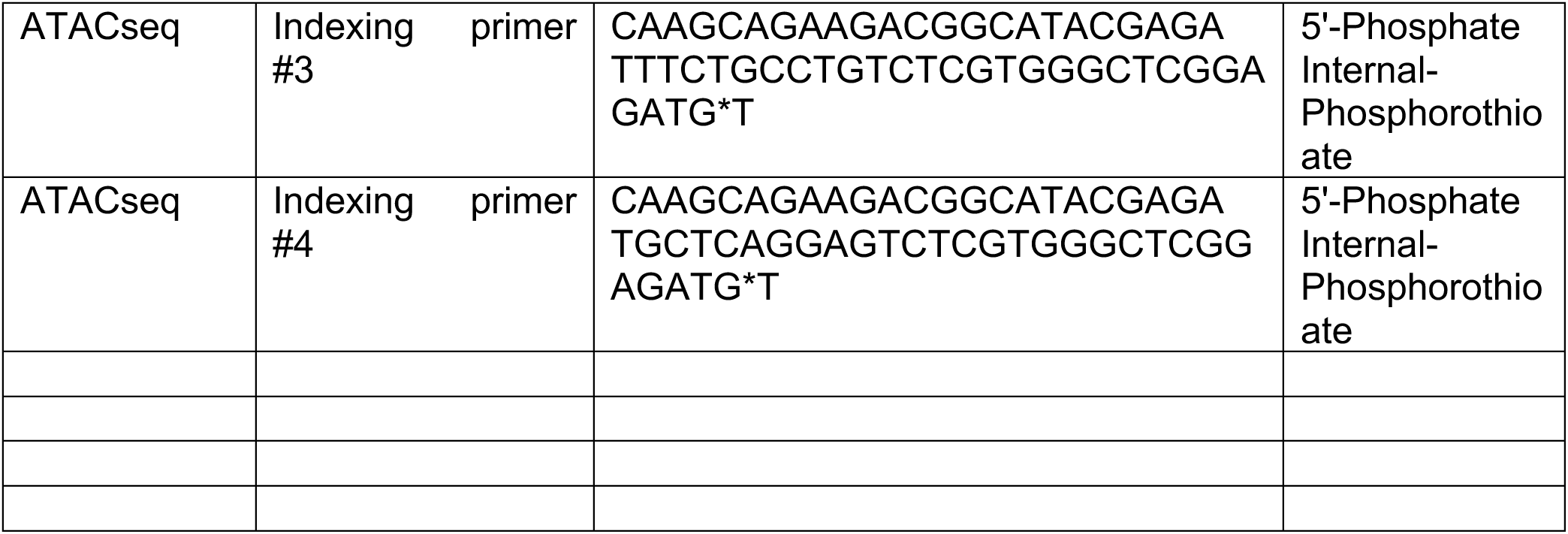

